# First intron length in mammals is associated with 5’ exon skipping rate

**DOI:** 10.1101/024463

**Authors:** Seung Gu Park, Sridhar Hannenhalli

## Abstract

The first introns in eukaryotes are much longer than downstream introns. While the functional roles of large first introns have been studied extensively, investigations into the mechanisms leading up to extreme lengths are limited. Prominently, Hong et al. noted that the first introns are predominantly in 5’ UTR and suggested that its lengthening may have resulted from a 5’-ward shifting of donor site due to a lower selection on splice site, as well as a selection to occlude upstream cryptic translation start sites. Here we suggest exon skipping as an alternative mechanism for first intron lengthening. Exon skipping results in consecutive introns becoming part of a single longer intron. We reasoned that a 5’-biased exon skipping rate could lead to longer introns toward the 5’-end of the gene, especially the first intron. Based on multiple datasets in human and mouse, we indeed found that internal exons toward the 5’-end of the gene are skipped significantly more frequently than the downstream exons. Importantly, we show that 5’-biased exon skipping is supported by consistent 5’-bias in several genomic, epigenomic, contextual, and evolutionary features that can be functionally linked to exon skipping. Interestingly, we found that first introns are enriched for relics of, now defunct, exons, some of which may have been recruited for regulatory functions; a significantly greater-than-expected fraction of such exons are included in cDNAs in other mammals. Overall, our results offer 5’-biased exon skipping as a novel, and arguably more potent, alternative explanation for substantially lengthening of first introns.

## Introduction

Genes in almost all eukaryotes have introns, which are spliced out in the mature mRNA. Considering the energetic costs of maintenance and transcription, the length and prevalence of introns seems puzzling, and have fueled much debate concerning their functional role (Berget et al. 1977; Chow et al. 2000; Duret 2001; Simpson et al. 2002; Fedorova and Fedorov 2005; Koonin 2006). Even more puzzling, several studies have revealed that in higher eukaryotes, especially in mammals, the first (5’-most) introns are much longer than downstream introns, especially when the first intron is within the 5’ UTR (Table 1). For instance, the first introns in human are on average more than 2-fold greater than the average length of all other introns (Bradnam and Korf 2008), and in extreme case it can be up to 1.5 Mb long. These observations have spurred investigations into the mechanisms underlying the length of first introns, and their functional consequences thereof (Table 1).

**Table 1.** List of studies related to first intron length.

| Main focus | Intron property | Data and Species | References |
| --- | --- | --- | --- |
| Longer Introns within 5'UTRs | Intron insertion | 2,057 introns (Vertebrates), 229 introns (Insecta), 126 (Fungi), and 200 introns (Plants) | (Hawkins 1988) |
|  | Splice-site shift to occlude uAUG | 10,562 transcripts (*Arabidopsis thaliana*), 3,424 transcripts (*Drosophila melanogaster*), 5,236 transcripts (Human), and 4,527 transcripts (Mouse) | (Hong et al. 2006) |
|  | AT-rich sequence in 5' UTR introns | 32,955 genes (*Arabidopsis thaliana*) | (Chung et al. 2006) |
| Longer first intron | Differential selection | 59 genes (from 8 species) | (Smith 1988) |
|  | Larger number of very long intron; In yeast, genes usually contain single intron | 367 genes (*Homo sapiens*), 140 genes (*Drosophila spp.*), 42 genes (*Zea Mais*), 139 genes (*Arabidopsis thaliana*), 73 genes (*Aspergillus spp.*), 101 genes (*Saccharomyces cerevisiae*), and 119 genes (*phylum Apicomplexa*) | (Kriventseva and Gelfand 1999) |
|  | To facilitate increased regulatory complexity | 9,499 genes (Human) | (Kalari et al. 2006) |
|  | Signal for transcription initiation and gene processing | 453 genes (Human Chr 21 and 22) | (Chen et al. 2002) |
|  | The higher divergence and higher proportion of functional elements | 7,791 orthologus (Human and Chimp) | (Gazave et al. 2007) |
|  | Enriched regulatory elements | 630 orthologus (*Drosophila melanogaster* and *Drosophila yakuba*) and 570 genes | (Marais et al. 2005) |
|  | Accumulation of functional intronic DNA | 6,381 orthologus (Mouse and Rat) | (Gaffney and Keightley 2006) |
|  | Enriched regulatory elements | 27,651 genes (*Arabidopsis thaliana*) | (Seoighe et al. 2005) |
|  | Enriched functional elements | 1,874 CDSs(*Arabidopsis thaliana*), 1,622 CDSs (*Caenorhabditis elegans*), 2,116 CDSs (*Drosophila melanogaster*), and 1,155 (Human) | (Bradnam and Korf 2008) |
|  | Enriched functional elements | 19,770 genes (Human) | (Li et al. 2012) |

In explaining the evolutionary maintenance of long first introns to overcome their energetic and maintenance costs, a vast majority of the previous studies have shown an enrichment of various putative regulatory signals in long first introns, consistent with a ‘genome design’ model (Vinogradov 2006). However, there have been very few studies specifically investigating the potential mechanisms leading to long first introns. Prominently, Hong et al. showed that introns within 5’ UTR are longer than introns within CDS (Hong et al. 2006). They suggested that this may be due to a combination of a lower selection on the precision of splice sites in the UTR, which can result in slight shifting of splice sites and an increase in intron length, as well as directional selection to occlude upstream cryptic translation start sites, again resulting in an increase in the intron length. While occlusion of upstream cryptic start codon can in principle be achieved by skipping of the exon, Hong et al. argued that this is not likely to be a major mechanism, because, in their dataset, 92% of 5’ UTRs span two or fewer exons, which precludes the possibility of exon skipping. Thus an increase in intron length, according to their model, can be primarily achieved by shifts of splice site. However, such a shift-based mechanism can mainly explain incremental increase in intron length. This previous analysis was based on then available full-length cDNA library.

However, in the recent human and mouse gene annotation datasets that we have compiled from two reference sources, when we consider all isoforms, we found that a substantial fraction of genes (48.7%) have more than 3 exons spanning the 5’ UTR, raising the possibility of exon skipping as a mechanism for long first introns. Moreover, it is difficult to explain very large first intron lengths by incremental shifts. Exon skipping, in contrast, can result in dramatic increase in intron length. Finally, 78% of the genes in our dataset have more than two alternative start codons (in alternative isoforms) with a mean distance of 10 kb between them, suggesting that various isoforms (from the same pre-mRNA) resolve among alternative start codons via exon skipping. Thus exon skipping, especially toward the 5’-end of the gene, may be responsible for very long first introns.

Exon skipping is one of the major categories of alternative splicing events, which is the main driver for generating functionally diverse proteins from limited number of genes (Chow et al. 1977; Berget et al. 2000; Pan et al. 2008; Barash et al. 2010; Florea et al. 2013). Several features associated with exon skipping have been reported, including, strength of splice sites, exonic splicing enhancers (ESEs) and silencers (ESSs), elongation rates of RNA polymerase II, nucleosome density, histone modifications, and binding of other splicing regulatory proteins (Table 2). Also, exon/intron architecture can influence the exon skipping event. For example, internal exons when truncated to less than 50 nucleotides tend to be skipped (Dominski and Kole 1991; De Conti et al. 2013). Also, internal exons when expanded to more than 300 bps can be skipped or have activation of cryptic splice site in it (Berget 1995; De Conti et al. 2013). Moreover, in Drosophila and human, exons flanked by long intron have a greater tendency to be skipped relative to those flanked by short introns (Fox-Walsh et al. 2005b; Roy et al. 2008a). All aforementioned studies have shown that length of exons and introns can influence exon skipping. While at the same time, skipped exons can manifest in long introns, essentially a composite of two adjacent introns; skipped exons within such composite introns have been experimentally demonstrated by lariat sequencing (Awan et al. 2013; Bitton et al. 2014).

**Table 2.**
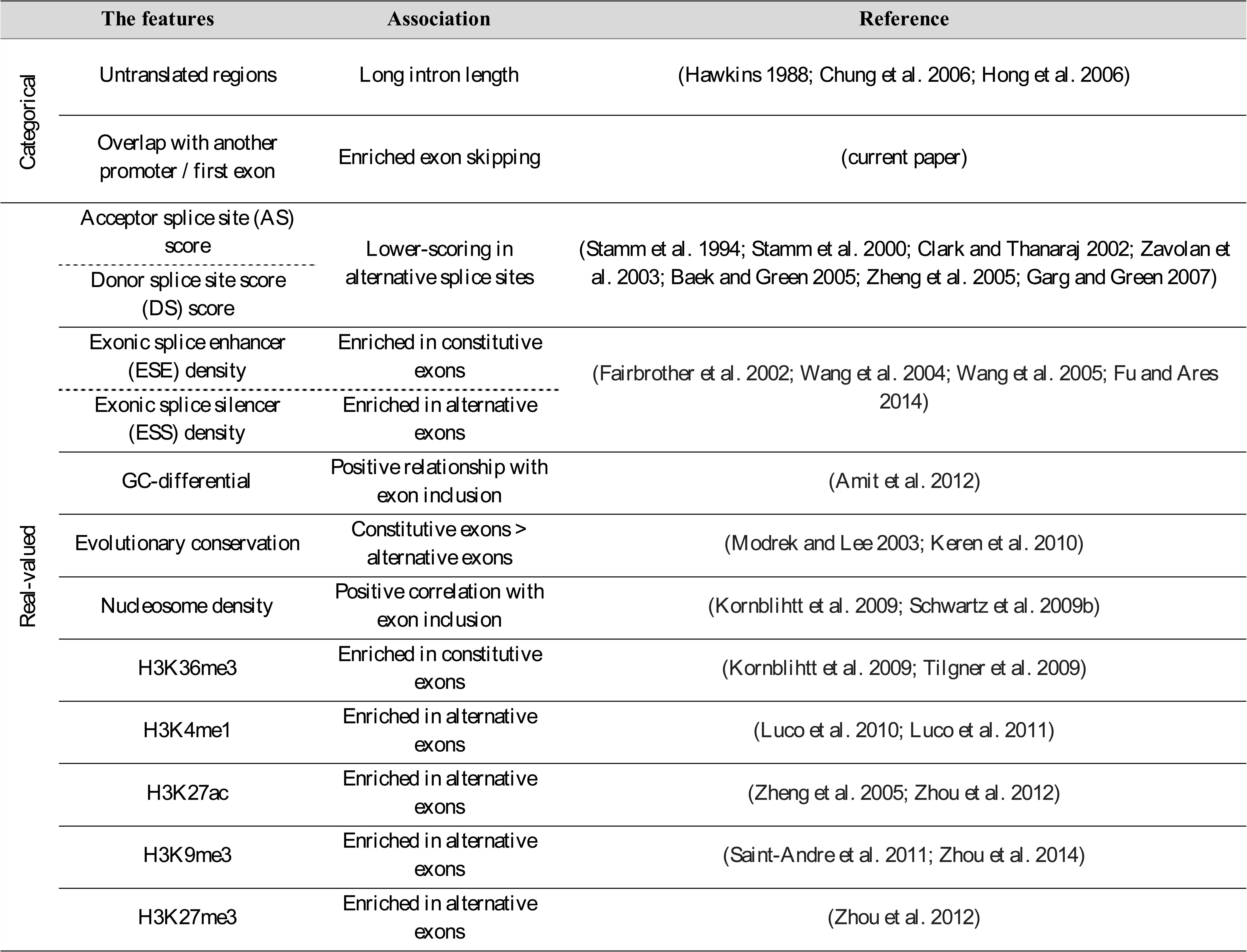
List of the features associated with alternative splicing or intron length.

Given that exon skipping can lead to long introns in the mature mRNA, in this work, we set out to assess the extent to which a greater length of first introns is explained by a greater exon skipping toward the 5’- end of the gene. Previous studies of intron lengths have considered individual mRNAs in isolation, however, we note that exon skipping event cannot be inferred when considering individual mRNAs in isolation, but only by jointly examining multiple isoforms. Therefore, we first constructed putative pre-mRNAs by grouping reference mRNAs of a gene based on identical 5’-end and for each such ‘pre-mRNA’ group, we quantified skipping events. We found that internal exons toward the 5’-end of the gene are skipped significantly more frequently, coincidental with long first introns. We further investigated several genomic, epigenomic, contextual, and evolutionary features that can be potentially functionally linked to exon skipping, including, UTR exon, overlap with other promoters and overlap with first exon, splice site scores, Exon Splice Enhancers (ESE), Exon Splice Silencers (ESS), GC-differential (difference between GC-content in an exon and flanking intronic regions), evolutionary conservation, nucleosome density, H3K36me3, H3K4me1, H3K27ac, H3K9me3 and H3K27me3 (Table 2). We ascertained that these features are associated with exon skipping rate and further show that these features also exhibit a strong bias toward 5’-end of the gene. Interestingly, we found that long first introns are enriched for putative internal exons, which may either be previously used now defunct, or rarely used (i.e., often skipped) exons. Moreover, we found such potentially pseudogenized exons to be enriched for transcriptional regulatory epigenomic signals, suggesting that some of them may be recruited for regulatory functions.

Overall, based on analyses in human and mouse, our results offer a novel alternative explanation for substantially greater lengths of first introns, namely, that longer first introns observed at the mRNA level are a manifestation of higher exon skipping rate toward the 5’-end at the pre-mRNA level, especially at the second exon, thus linking the inordinately long first introns in eukaryotes with alternative splicing and exon turnover.

## Results

### A pre-mRNA view of gene transcripts

Introns are defined based on the mapping between mature mRNA and the genome. However, splicing acts on the pre-mRNA. Thus based on the mRNA definition, a long intron may result from exon skipping events at the pre-mRNA (pre-splicing) stage. However, a direct observation of exon skipping events at the pre-mRNA level at high throughput is currently not feasible (Awan et al. 2013; Bitton et al. 2014). Therefore, we first devised an approach to infer exon-intron structure, and thus potential exon skipping events, at the pre-mRNA level by integrating mRNA data (Fig. 1). Briefly, for each gene, we grouped mRNAs based on their start site locations, and identified the largest such group mRNAs. We then constructed *a pre-mRNA template* where the first and the last exon of the pre-mRNA template is formed by collapsing respectively the first and the last exons of all the mRNA in the group, and the internal exons of the pre-mRNA template are formed by collapsing all exons from all mRNAs for the gene, into meta-exons. Thus in the pre-mRNA template the introns do not overlap an internal exon for any mRNA; we refer to the introns at the pre-mRNA level as *pure introns* (Fig. 1).

**Figure 1.**
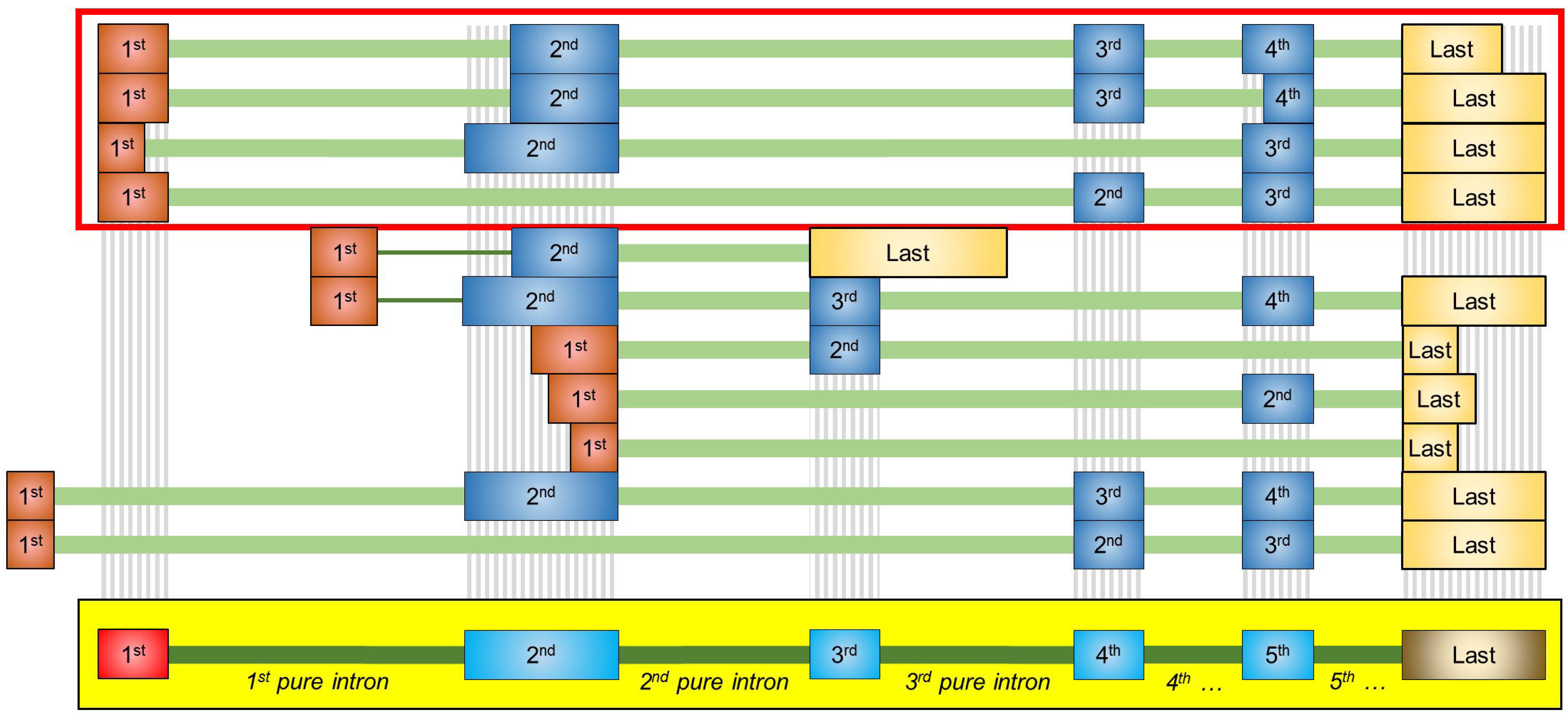
Construction of putative pre-mRNA template. Each row represents a transcript, with exons shown in rectangles and intron with think green line. Exon ordinal positions are shown in exon boxes. Dark orange boxes represent first exons. Internal exons are shown in blue. Starting with all mRNAs for a gene, largest set of transcripts with identical start locations was identified as the representative pre-mRNA group (Red box in the top). First exons in the pre-mRNA group are collapsed to obtain the first meta-exon (red rectangle) of the meta-transcript or the pre-mRNA template (yellow box). All internal exons are collapsed to form internal meta-exons (light blue rectangles) of pre-mRNA template. Last exons (light gold rectangles) of pre-mRNA group are merged into last (gold rectangle) meta-exon of the pre-mRNA template.

We obtained exon-intron annotations for transcripts from Ensembl (Cunningham et al. 2015) and UCSC table browser (Karolchik et al. 2004) and obtained the intron lengths. As noted previously, we found that the first (5’-most) introns are significantly longer than downstream introns (Fig. 2). We found that first pure introns are also longer than downstream pure introns (Fig. 2), but as expected, pure introns (pre-mRNA view) are significantly shorter than the introns (mRNA view). Interestingly, however, this difference in intron lengths between the mRNA and the pre-mRNA view was found only for the first (*p* = 4.91e-09) and to a lesser extent for the second (*p*=2.98e-02) intron, and not for downstream introns (Fig. 2). We observed a similar trend in mouse (Supplementary fig. 1). These results are consistent with a higher rate of exon skipping at the 5’-end, thus resulting in shorter pure introns (relative to introns in mRNA view) specifically at the 5’-end of the gene.

**Figure 2.**
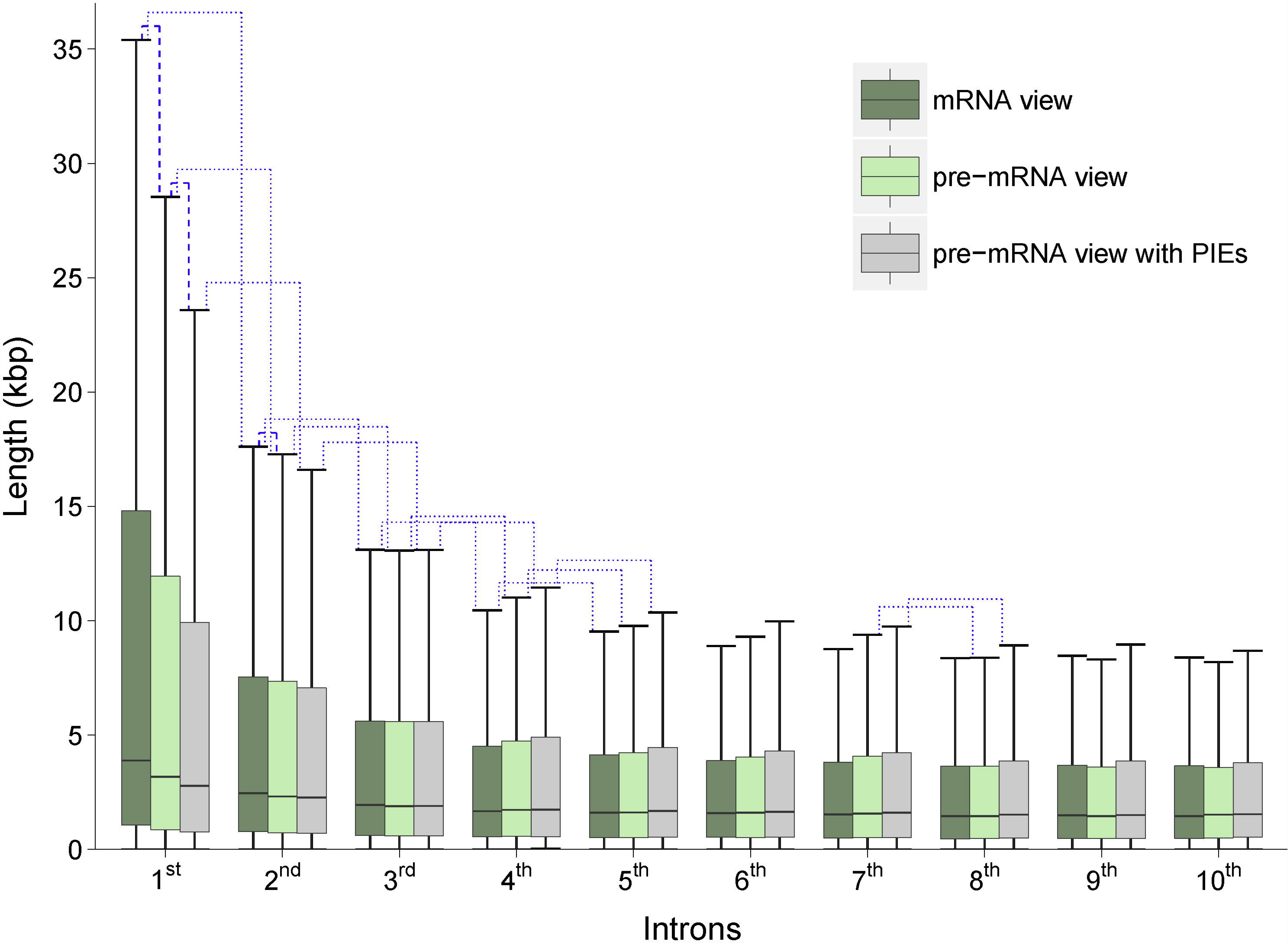
Human intron length comparison for different views of introns. Dark green: conventionally calculated intron lengths from all mRNAs. Light green: “*pure intron*” lengths from our constructed pre-mRNA templates. Gray: the intron lengths from pre-mRNA view when PIEs are included (see text). Both dotted lines linked between ordinal positions within the group and dashed line linked between groups within ordinal position represent significant *p*-values estimated by one-sided Wilcoxon rank sum tests between linked groups, blue color is computed with one-sided “greater than” the alternative hypothesis and red color show the opposite alternative hypothesis, “less than”. All results are obtained from human, and the results from the mouse are provided in Supplementary fig. 1.

### Five prime exons are skipped more frequently than downstream exons

Next we tested the hypothesis that a significant length difference in first and second intron between mRNA definition and pre-mRNA definition comes from frequent exon skipping for 5’-exons. First exons are modified by 7-methylguanosine cap and are integral part of the mRNA as they play a critical role in splicing of downstream exons, and are thus immune to skipping (Izaurralde et al. 1994; Berget 1995; De Conti et al. 2013; Kornblihtt et al. 2013). We therefore analyzed the skipping rates of internal exons only. Previous studies have shown that short (<50 bp) and long (>300 bp) internal exons tend to be skipped (Dominski and Kole 1991; Berget 1995; Fox-Walsh et al. 2005a; Roy et al. 2008b; De Conti et al. 2013). As shown in Supplementary fig. 2, we found that the second and third exons (i.e., first and second internal exons respectively) indeed have a relatively large proportion of short and long exons. Thus the unusual exon length in the 5’-end may result in their higher skipping rate. However, to investigate additional biological features potentially affecting exon skipping at the 5’-end, in what follows, we excluded the short and long exons from all analyses.

The introns in higher eukaryotes are on average 21-fold longer than exons. Therefore during splicing, coordinating the interactions between the two ends of an intron is substantially more challenging and error prone relative to coordinating interactions between the two ends of an exon. This realization has led to the ‘*Exon definition*’ model of splicing (Robberson et al. 1990; Berget 1995; De Conti et al. 2013). However, we noticed that exons with short flanking introns tend not to get skipped compared to exons with long flanking introns (Supplementary fig. 3). Because this suppression of exon skipping preferentially affects downstream exons due to short flanking introns, this may inflate the relative exon skipping rate for the 5’-exons. To remove this bias, we therefore excluded from our analyses the exons that are flanked by short introns (< 300bp). However, we note that exons with short flanking introns have similar relationship between skipping rate and ordinal position as for exons with long flanking introns (Supplementary fig. 3).

After these cautionary filtering of the data, we estimated exon skipping rate for each meta-exon based on the presence/absence of meta-exon nucleotides in different mRNAs in a pre-mRNA group; we will refer to this set of estimated exon skipping rates as the “Reference dataset”. To ascertain the robustness of our results we utilized two additional sources for exon skipping rates: (1) HEXEvent database (Busch and Hertel 2013), which estimates exon skipping rates from EST libraries, and (2) exon skipping events from MISO annotation dataset based on multiple RNA-seq libraries (Katz et al. 2010). As shown in Fig. 3, in all three datasets, internal exons toward 5’ are skipped at a higher frequency, especially the second and the third exons. We found a similar trend in mouse using both the reference and the MISO dataset (Supplementary fig. 4). Overall, these results establish a greater rate of exon skipping toward the 5’-end, which can potentially explain longer first introns to some extent.

**Figure 3.**
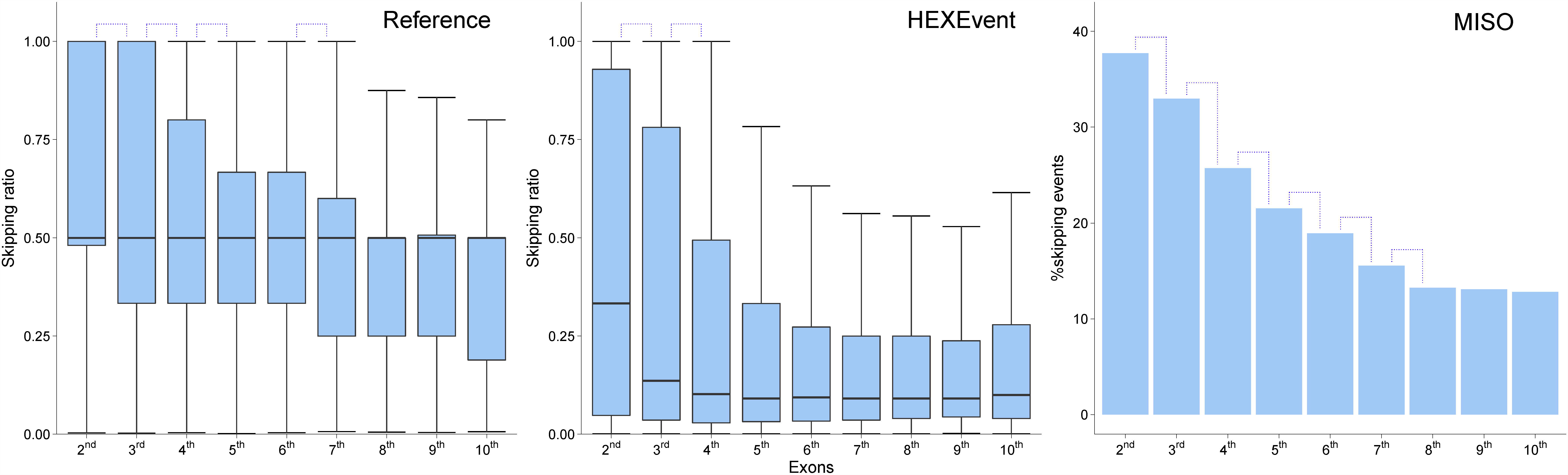
Skipping rates for internal exons in human. The three panels correspond to three different sources used to compute exon skipping rates of internal exons. The exon skipping rates from “Reference” and the “HEXevent” datasets are box-plotted which have one-sided Wilcoxon rank sum test between “N^th^” exons and “(N+1)^th^”. For “MISO” dataset the proportion of skipped exon are bar-plotted, and one-sided Fisher’s exact tests are performed. Significant *p-*values are represented by blue dotted line. The odds ratios for significant *p-*values on MISO datasets from the left dotted lines are 1.2, 1.4, 1.2, 1.2, 1.3, and 1.2 respectively. All results are obtained from the human dataset, and the results from the mouse are provided in Supplementary fig. 4.

### Five prime biased exon skipping is associated with several genomic, epigenomic, and contextual features

Next, for a number of genomic, epigenomic, and contextual features relevant to splicing, we investigated their association with the observed greater rate of exon skipping. We considered the following features (see Methods, Table 2): (1) whether or not the exon is untranslated (5’ and 3’ UTR), (2) whether the exon overlaps a promoter (2kb upstream of the first exon) corresponding to another pre-mRNA, (3) whether the exon overlaps a first exon corresponding to another mRNA, (4) Acceptor splice site (AS) score, (5) Donor splice site (DS) score, (6) Exonic Splice Enhancer (ESE) density, (7) Exonic Splicing Silencer (ESS) density, (8) GC differential, (9) Exon conservation, (10) Nucleosome density, (11) H3K36me3, (12) H3K4me1, (13) H3K27ac, (14) H3K9me3, and (15) H3K27me3. The first three of these are categorical features, and the rest are real-valued features. Using both the reference and HEXEvent datasets, we first assessed the associations between skipping rate and various features, regardless of the ordinal position of the exon.

As shown in Table 3A, we found that most features exhibited significant association with exon skipping rate. In all but one case (H3K27me3) when the results are inconsistent, result in only one of the two directions is significant. Broadly, among the continuous features, AS score, DS score, ESE, GC-differential, Exon conservation, nucleosome density and H3K36me3 show a significant negative correlation with exon skipping rates. ESS, H3K4me1, H3K27ac, H3K9me3 and H3K27me3 on the other hand, shows a significant positive correlation with exon skipping. For the four categorical features, being UTR or overlap with alternative promoter and first exon are associated with significantly higher skipping rates while coding for a protein, as expected, has the opposite association (Table 3B).

**Table 3A.**
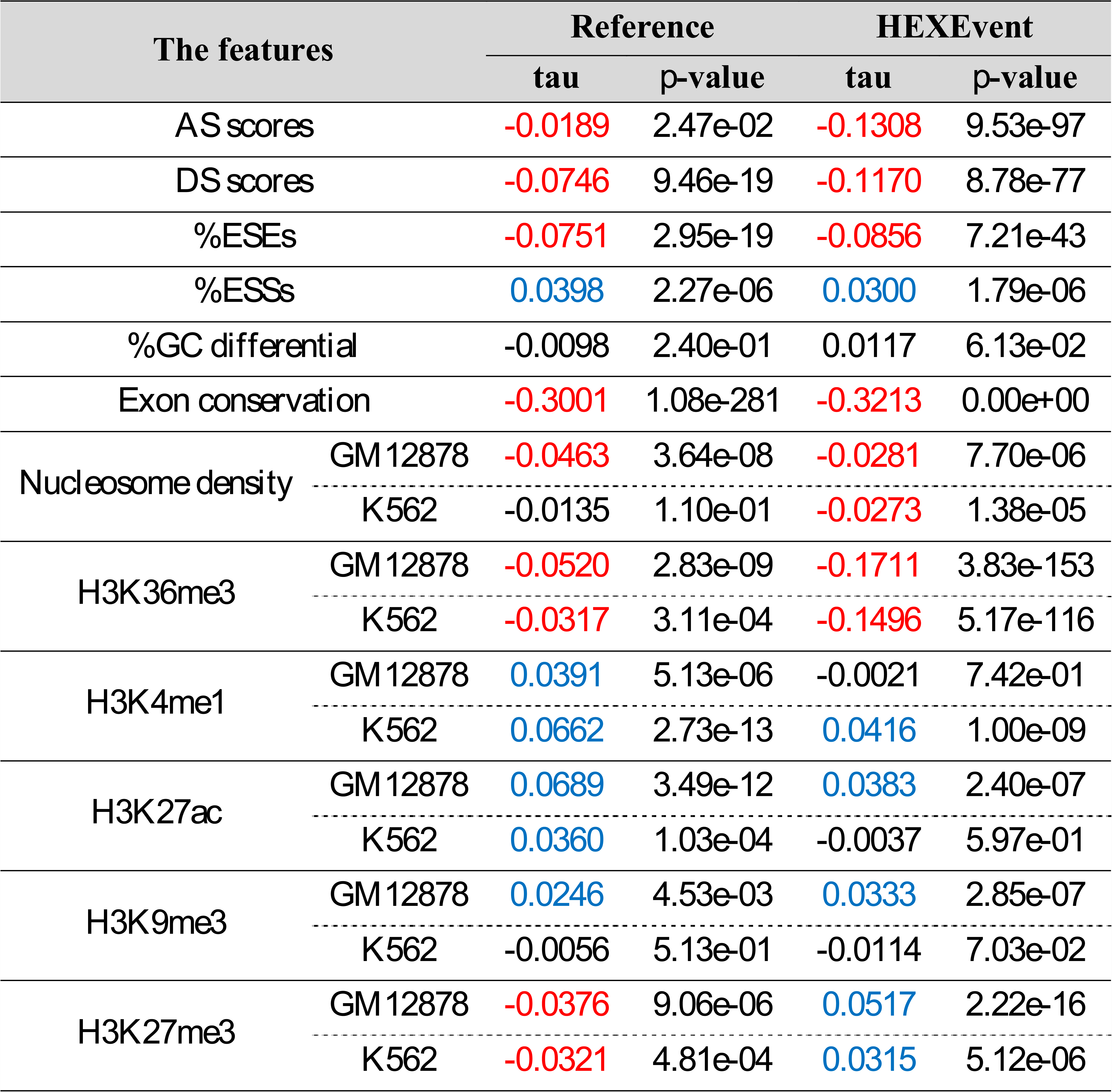
Correlation test between exon skipping rates and the features.

**Table 3B.**
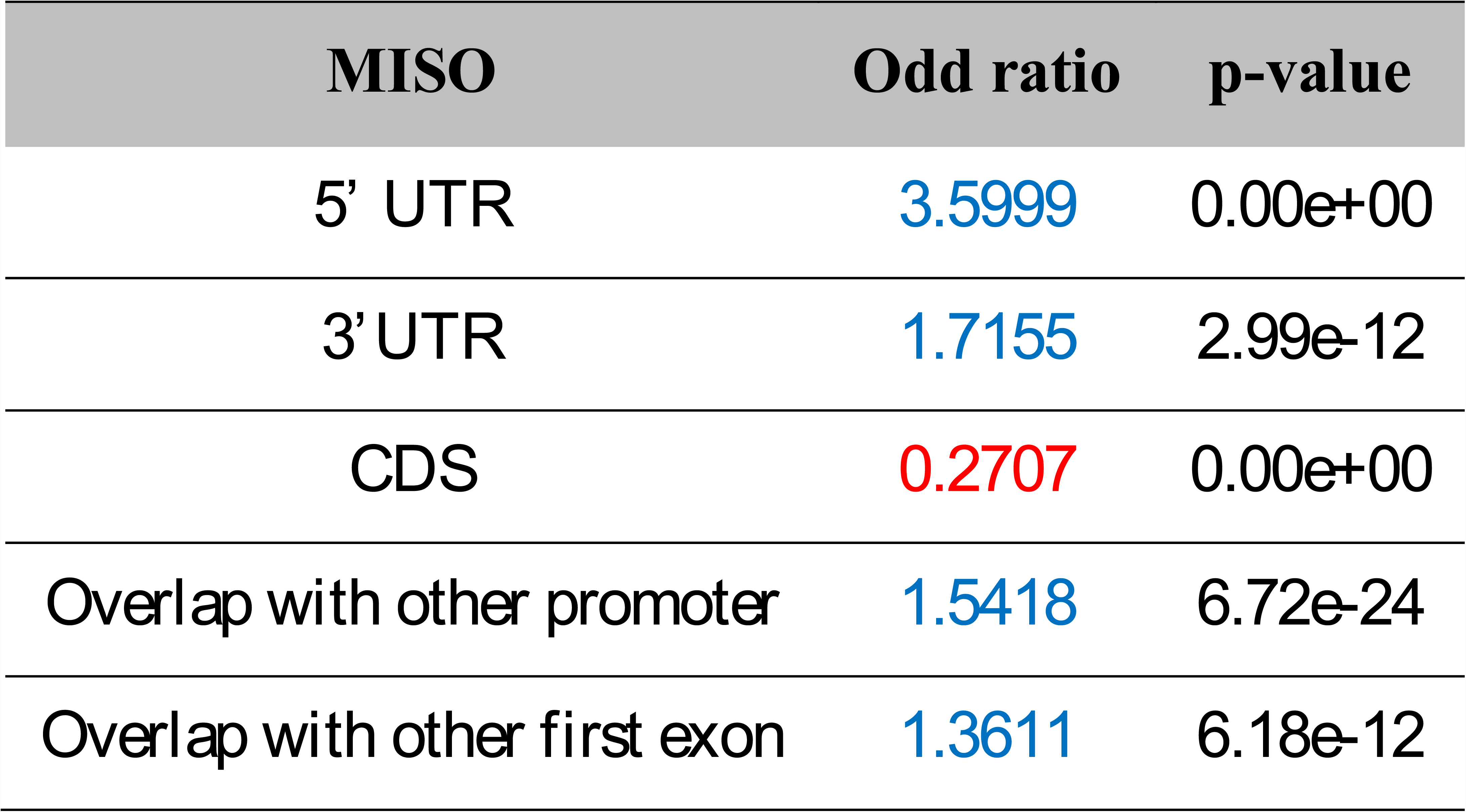

In addition to the association test, we analyzed the trend of feature value by quantile of exon skipping rates (for continuous features), or skipping/non-skipping ratio (for categorical features using MISO). Supplementary fig. 5 shows these trends are somewhat consistent with the association analyses. But in the case of AS score and GC differential, in some cases the trend analysis is not consistent in direction with the association test.

Next we assessed whether the features that are associated with exon skipping also exhibit 5’-bias, which may provide clues to potential mechanisms underlying the observed 5’-bias of exon skipping. As shown in Fig. 4, for most features (except for AS and DS score), we see a significant 5’-bias; in particular there is a significant difference between the second and the third exon. An enrichment of 5’ UTR at the second exon is expected. While a greater tendency for the second exon to overlap with an alternative promoter and first exons of other transcripts is also unsurprising, this overlap of function can nevertheless be mechanistically linked to weaker splicing signals and therefore greater skipping of such exons. Previous studies have reported lower evolutionary conservation at the 5’ UTR exons than CDS (Shabalina et al. 2004). High proportion of 5’ UTR at the second exon is thus expected to result in less conserved second exons. We compared evolutionary conservation across meta-exons, using PhastCons score based on 100 vertebrate species (Siepel et al. 2005). As shown in Fig. 4, the second exon and its flanking region were significantly less conserved than downstream exons. Interestingly, even though there is a peak of conservation at GT/AG consensus sequences at exon-intron junction (known to be critical for splice site recognition (Mount 1982; Burset et al. 2000)), the absolute value of the conservation (and therefore, presumably, the selective constraint) is significantly lower for the second exon. Also, the difference in conservation between second and downstream exons disappears sharply beyond the splice sites. These results are also consistent when we used PhastCons score based on 46 mammalian species (Siepel et al. 2005)(data not shown).

**Figure 4.**
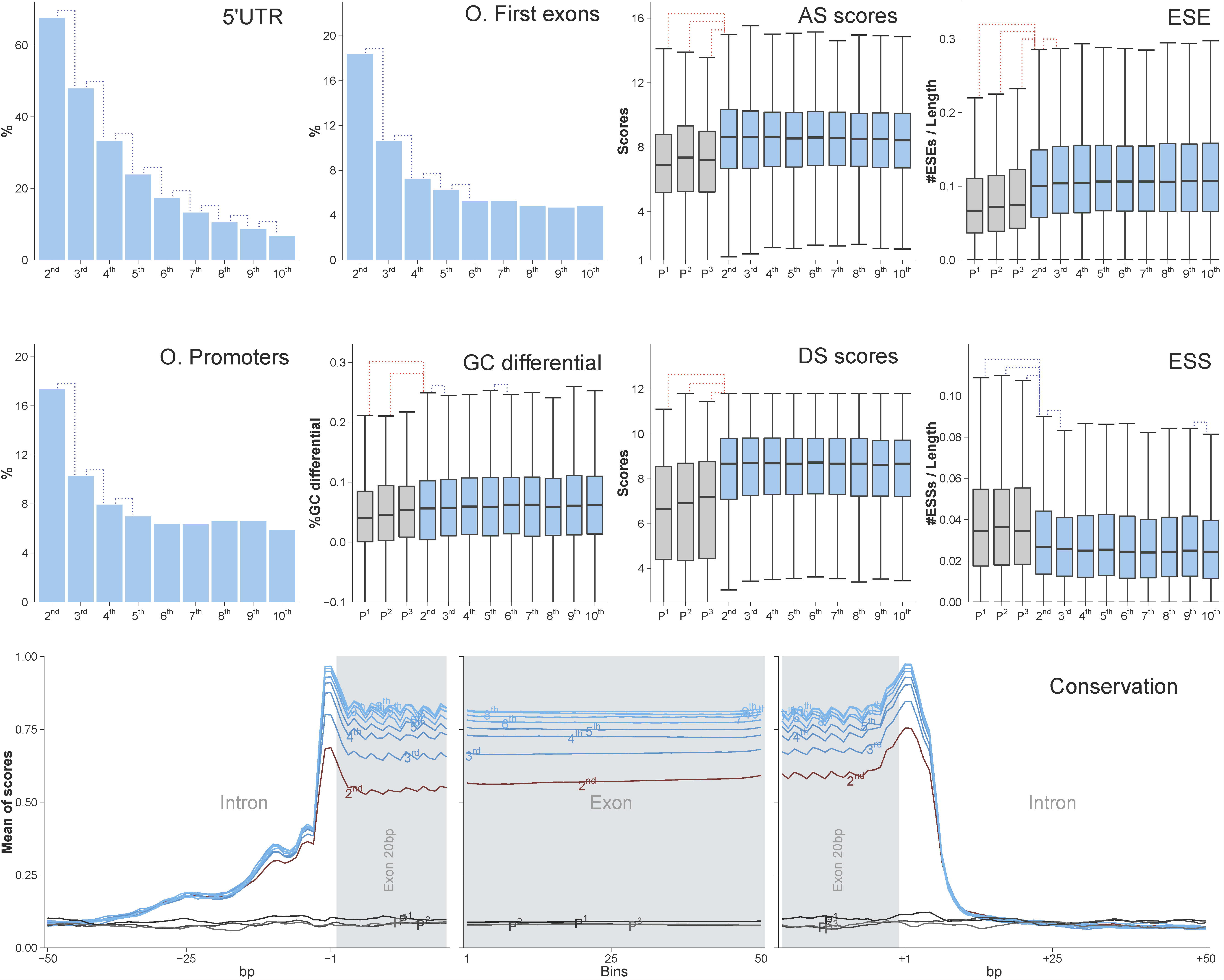
Comparison of various genomic and functional features between exons. Investigated feature is shown in right-top of each plot. The proportion of 5’ UTR exon, proportion of overlapped exon with other promoter and other first exon are bar-plotted. GC differential, AS scores, DS scores, ESE and ESS are box-plotted. Blue bars and boxes represent 2^nd^ to 10^th^ exons. Gray boxes represent PIEs in pure intron. All dotted lines show significant *p*-values estimated by one-sided Wilcoxon rank sum tests between linked groups, blue color is computed with one-sided “greater than” alternative hypothesis and red color show the opposite “less than” alternative hypothesis. In the bottom row, mean of phastCons scores on nucleotides or bins are plotted. Gray box indicates exon region (exons were averaged to fit in 100 bins). The 50bp flanking regions of exon box are intron regions including 20bp exon. The ordinal position of exons are shown on the line, especially second exon are shown red line. And gray lines indicate PIEs in pure introns. The odds ratios for significant *p-*values on the 5’ UTR datasets from the left dotted lines are 2.2, 1.8, 1.6, 1.5, 1.4, 1.3, 1.2 and 1.3, and on the overlap with other promoter (O. promoters) group are 1.8, 1.3 and 1.2, and on the overlap with other first exons (O. first exons) group are 1.9, 1.5, 1.2 and 1.2.

Because promoter, 5’ UTR and first exon have higher GC-content than internal exons (Zhang et al. 2004), their overlap with second exons leads to the enrichment of GC-content at second exons; medians of GC-contents at second exons and at third exons are 0.4862 and 0.4783 respectively (*p*=6.16e-07). Fig. 4 also shows a clear 5’-bias for splicing signals such as ESS, and ESE; while ESE density is lower at the second exon, ESS density shows the opposite trend, consistent with their repressive role. ESE hexamers are known to be enriched for adenine (47%, compared to the genomic background 30%) (Fairbrother et al. 2002; Fairbrother et al. 2004). Consistently, we see a strong negative correlation between GC-content and proportion of ESE (Kendall’s tau=-0.3606, *p*=0.00e+00). These compositional properties can thus explain lower-ESE in second exon, which has high GC-content. However, ESS have weak negative correlation with GC-content (Kendall’s tau=-0.0162, *p*=5.50e-09).

Overall, we see a strong and consistent trend for a 5’-bias in the features that correlate with exon skipping, mechanistically linking them to the greater skipping rate for the 5’ internal exons. Interestingly, AS and DS score do not exhibit a 5’-bias, despite the fact that they show a clear correlation with exon skipping rate in general (Stamm et al. 1994; Stamm et al. 2000; Clark and Thanaraj 2002; Zavolan et al. 2003; Baek and Green 2005; Zheng et al. 2005; Garg and Green 2007); we discuss this later.

### Epigenomic signals at second exons are associated with exon skipping

We found that the nucleosome occupancy scores at second exons are significantly lower than those at the downstream exons, across the exon region as well as the flanking regions (Fig. 5 and Supplementary fig. 6). These is consistent with previous reports suggesting that the nucleosome occupancy of low inclusion exons is lower than high inclusion and constitutive exons (Schwartz et al. 2009a). Nucleosome occupancy is also related to GC-differential between exon and flanking intron. Even though second exons have high GC-content, due to high GC-content at first introns (first intron=0.4707, second intron=0.4352 and *p*=2.54e-102), as shown in Fig. 4, ‘GC differential’ at second exons is lower than downstream exons. Low GC-differential at second exon results in lower nucleosome density resulting in a greater transcription elongation rate, which is known to promote exon skipping (Amit et al. 2012).

**Figure 5.**
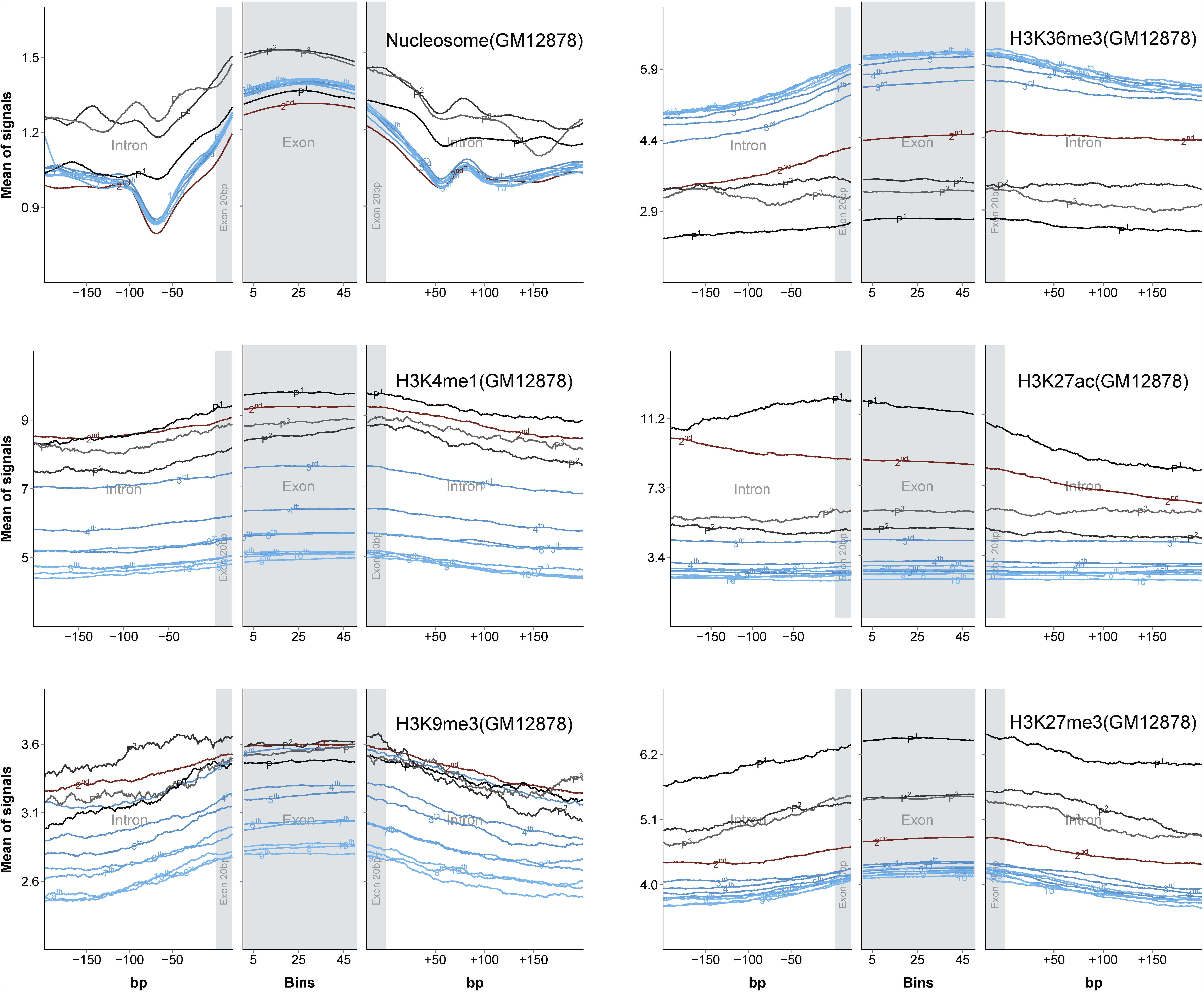
Comparison of epigenomic features between exons. Each epigenomic signal was mapped to exon and their 200 bps flanking introns including 20bp exon. Bright gray box indicate exon region in which exons were averaged to fit 100 bins. The ordinal position of exons are shown on the line, second exon are shown in red and downstream exons are shown blue line. Gray lines indicate PIEs in pure introns. All results are obtained from the GM12878 cell line, and the results from the K562 cell line are provided in Supplementary fig. 6.

Interestingly, several histone marks known to be associated with transcriptional processes exhibit differential occupancy at second exons. H3K36me3 marks – which are related to transcription elongation and are typically found in constitutive exons – are substantially lower, consistent with faster elongation, at the second exons than those at the downstream exons (Bannister et al. 2005; Vakoc et al. 2006; Barski et al. 2007; Kolasinska-Zwierz et al. 2009; Tilgner et al. 2009; Kornblihtt et al. 2013). Other histone modifications – H3K4me1, H3K27ac, H3K9me3, and H3K27me3, associated with transcriptional regulatory functions, are higher at second exons than at downstream exons as well as their flanking regions (Barski et al. 2007; Benevolenskaya 2007; Heintzman et al. 2009).

To ensure that observed trends related to second exons are not simply due to their proximity to transcription start site, we investigated the enrichment trends for all epigenomic marks after controlling for their distance for the 5’-end of the gene. We only used exons that were 1,000∼ 10,000bp away from the 5’-end of their gene. As shown in Supplementary fig. 7, the relative values of the epigenomic signals of second exons show similar trends, despite being based on small number of exons. Similarly, to ensure that the observed trends are not due to overlaps with alternative promoters, we also repeated the analyses after excluding the exons that overlapped an alternative promoter, and observed similar trends (Supplementary fig. 8 and 9). In this smaller group of exons, compared with all exons, the mean signal density is reduced only for H3K4me1 (*p*=2.45e-02) and H3K27ac (*p*=7.92e-03) which are associated with enhancers and promoters respectively (Barski et al. 2007; Benevolenskaya 2007) (Supplementary fig. 9), suggesting that second exons often overlap an alternative promoter. Given the cell type specificity of the epigenomic signal, we ensured that the above trends, which were performed in GM12878 cell type, are consistent in K562 cell lines (Supplementary fig. 6-9).

Overall, these analyses firmly establish a link between differential density of various epigenomic marks at 5’ internal exons and their higher skipping rates.

### Abundance of putative exons in first introns

Collectively, the results above suggest an overall lower efficiency of processing exons toward the 5’-end, which, we conjecture, results in a greater rate of exon skipping. Furthermore, a lower conservation at 5’ exon and broadly at exons with higher skipping rate suggests a relaxed functional constraint at these exons, which may lead to a higher rate of pseudogenization of exons at the 5’-end of the gene, resulting in constitutive longer first introns. This model predicts a greater density of pseudo-exons in first pure introns. Also in general we expect pseudo-exons to have features consistent with lower splicing efficiency and lower probability of inclusion in a mature product.

As a proxy for pseudo-exons, we used GENSCAN-predicted (Burge and Karlin 1997) putative internal exons (PIEs) that do not overlap any annotated exons and non-coding transcripts in the database.

We identified PIEs in the first, second, third, fourth and the fifth pure introns and compared the number of PIEs in introns after controlling for intron length. Supplementary Fig. 10 shows the sampled intron lengths. As shown in Fig. 6, first and second introns are significantly enriched for PIEs relative to second and third intron respectively (odds ratio=1.51, *p*=1.96e-04; odd ratio=1.30, *p*=2.95e-02).

**Figure 6.**
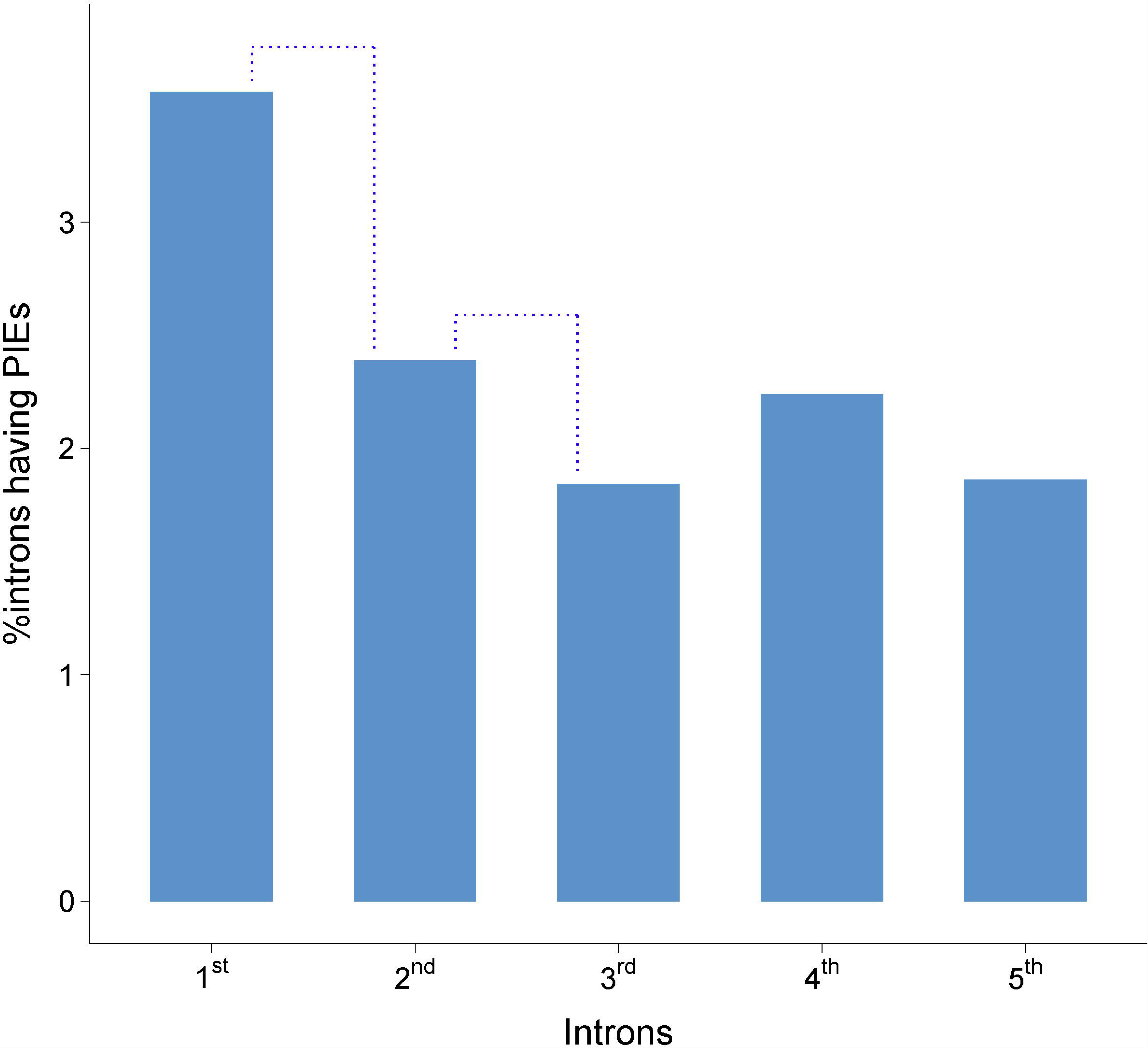
Proportion of intron having predicted internal exons in pure introns. The proportions of introns having PIEs are calculated from the sampled pure introns (Supplementary fig. 10). The blue dotted lines indicate significant *p-*values. The odds ratios for significant *p-*values is 1.5. The *p*-values are estimated by one-sided Fisher’s exact test between linked groups.

Next, similar to the characterization of reference exons above, we analyzed the properties of the PIEs; P^1^, P^2^, P^3^ refer to PIEs in the first, second, and the third pure intron respectively, in terms of AS score, DS score, ESE, ESS, GC-differential, evolutionary conservation and epigenomic marks. As shown in Fig. 4 (gray boxplots or gray line), consistent with our proposed model, splice site scores of PIEs are lower than those for the reference exons. Also proportion of ESEs is lower in PIEs than in second exons (Fig. 4); this is also true when PIEs were compared against downstream reference exons (Fig. 4). Consistent with negative correlation between GC-content and ESE density, GC-content of PIEs (in the first, second and the third pure intron) is higher than that for reference exons (median: reference exons=0.4734, PIEs=0.5321, *p*=1.05e-223); Interestingly, even though ESS have weak negative correlation with GC-content, PIEs have higher proportion of ESSs, suggestive of directional evolution.

Note that GC-differential of second exon is lower than the downstream exons. We found that GC-differential of PIEs in first and second pure introns are lower than those of reference exons (Fig. 4), consistent with higher skipping rate. Consistently PIEs and their flanking regions are also under a lower evolutionary constraint (Fig. 4). Moreover, H3K36me3 signals, known to associate with included exons, are also lower at PIEs than at reference exons (Fig. 5).

The results above collectively indicate lower efficiency for inclusion and lower functional constraint at PIEs consistent with their pseudogenization. Previous studies have shown an enrichment of transcriptional regulatory epigenomic signals in long first introns (Park et al. 2014). We therefore analyzed epigenomic signals relevant to transcriptional activation and repression in the context of PIEs. Interestingly, we found that the nucleosome density and epigenomic signals indicative of transcriptional repression, *viz*., H3K9me3 and H3K27me3 in GM12878 cell line, and H3K27me3 in K562 at the PIEs and flanking region are higher than not only the reference second exons, it is also higher than downstream exons. In contrast, the activation signals H3K4me1 and H3K27ac are lower than second exon and higher than downstream exons, with the exception of H3K27ac in GM12878 (Fig. 5 and supplementary fig. 6). This result may suggest that PIE may have acquired regulatory functions.

It is possible that PIEs represent rarely included exons in conditions that are not represented in the current databases. We therefore assessed the effect on intron length if all PIEs were included in rare conditions and were therefore in the reference dataset. We mapped all PIEs onto putative pre-mRNAs and recalculated the lengths of pure introns. As can be seen in Fig. 2, as expected, the lengths of re-built pure first intron are significantly shorter than those of putative pre-mRNA view.

### Evolutionary support for functionality of human PIEs in mammals

Our results thus far suggest that PIEs in human first introns may be a product of pseudogenization of exon of once functional exons in human. We therefore assessed whether some of the human PIEs may still be functional in other species. While true functionality is difficult to ascertain, we assessed whether orthologous sequences in other species are incorporated in a full length cDNA in the respective species. Using BLAST, we searched for human PIEs in the first introns against cDNA sequences in 37 mammalian species (Supplementary Fig. 11). At various BLAST E-value threshold, the numbers of BLAST hits against each species, as expected, is positively correlated with the evolutionary distance of the species from human (Kendall’s tau= 0.50-0.54, *p*=3.3e-06 - 1.8e-05). As a control for PIEs, we used randomly selected length-matched regions from the first pure introns in human; this controls for aberrant inclusion of intronic sequences into cDNA. For various E-value thresholds, we estimated the odds-ratio for the fraction of hits for PIEs and the controls, as well as the p-values for enriched hits of PIEs in mammalian cDNAs. Even though the absolute fraction of PIEs with an unambiguous match in another species is small (due to divergent evolutionary histories of gene structure evolution during mammalian evolution (Subramanian and Kumar 2003; Lorente-Galdos et al. 2013)), the PIEs are significantly more conserved in different species, at a wide range of BLAST thresholds (Fig. 7).

**Figure 7.**
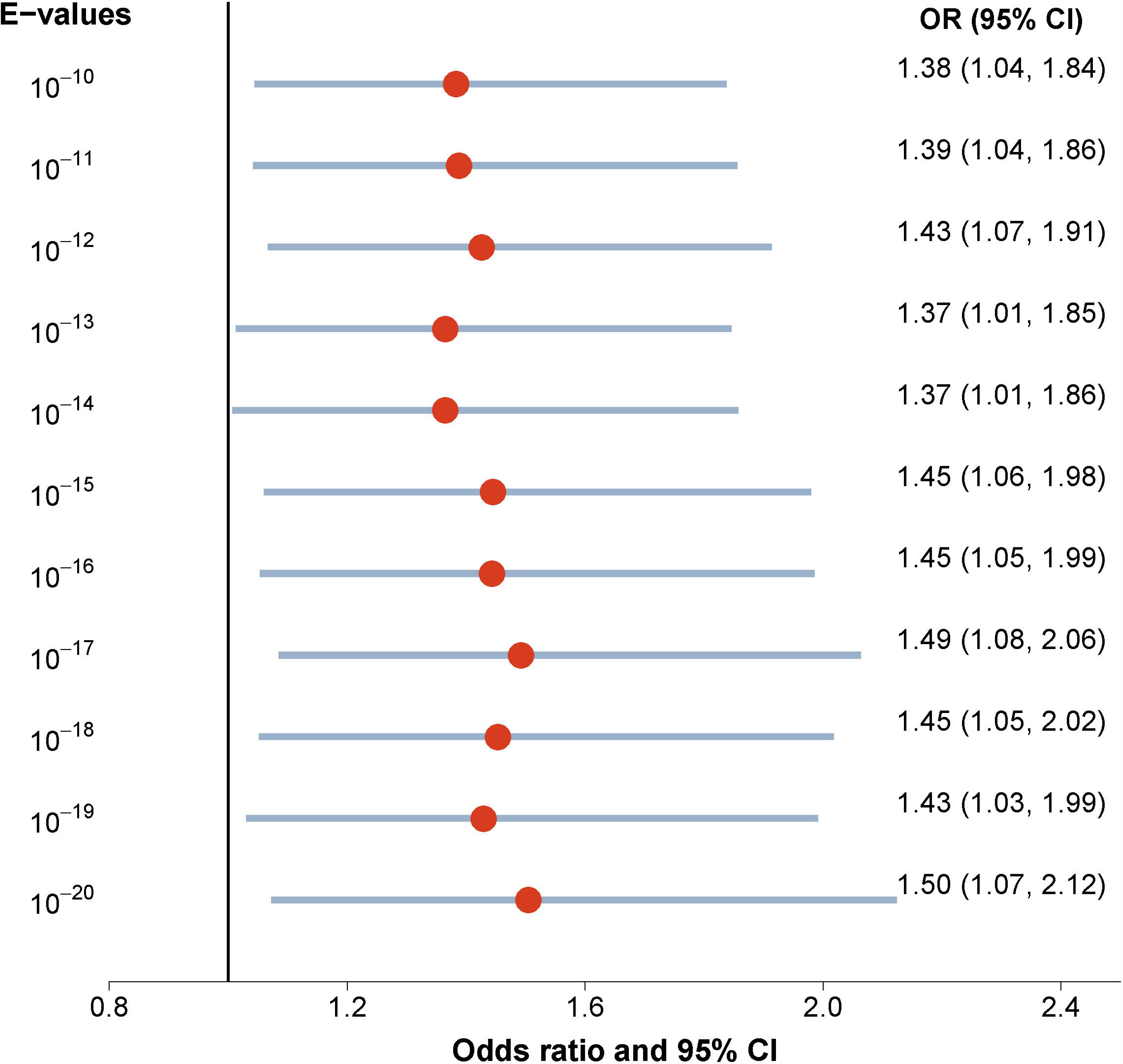
The odds ratio (OR) and 95% confidence intervals (CI) for the number of BLAST hits of PIEs. BLAST search of PIEs and randomly generated PIEs are performed against cDNA libraries in 37 mammals. Using the number of BLAST hits/non-hit of PIEs and those of randomly generated PIEs, OR and CI were calculated for different BLAST E-value cutoff (10^-20^ to10^-10^). Red circles and light blue bar indicates OR and 95% CI respectively. Exact numbers of hits on different E-values cutoff in 37 mammals are presented in Supplementary Fig. 11.

### Discussion

Existence of introns has intrigued researchers since their discovery (Gilbert 1978), considering not only the energetic costs of their maintenance, but also the requisite accuracy in splicing to generate the correct product (Berget et al. 1977; Chow et al. 2000; Duret 2001; Simpson et al. 2002; Fedorova and Fedorov 2005; Koonin 2006). The most significant advantage of having introns, counterbalancing the costs, is the fact that introns enable alternative splicing which generates the vast variety of isoforms, and also facilitate modular combination of protein domains in some cases (Brinster et al. 1988; Le Hir et al. 2003; Lynch and Kewalramani 2003; Wang et al. 2007). Parallel to the debate over existence of introns in the first place, is the debate over the prevalence of unusually long first introns in higher eukaryotes from *Coprinopsis cinerea* to *Homo sapiens* (Bradnam and Korf 2008). While functional implications of long first introns have been reasonably well studied (Smith 1988; Kriventseva and Gelfand 1999; Chen et al. 2002; Marais et al. 2005; Seoighe et al. 2005; Gaffney and Keightley 2006; Kalari et al. 2006; Gazave et al. 2007; Bradnam and Korf 2008; Li et al. 2012; Park et al. 2014), studies focusing on the genesis of long introns and our understanding of the mechanisms underlying the lengthening of first intron is limited (Hong et al. 2006) (Table 1).

One previous work has directly addressed the potential mechanisms resulting in the lengthening of the first intron over time. Hong et al. (2006) suggested that being predominantly UTR, there is lower selection on splice site conservation at first introns, allowing for shifting of splice sites at these introns. Leveraging the relaxed selection, first introns lengthen by shifting of splice sites under a directional selection to occlude cryptic upstream start codons and the same force acts against the shortening of these introns. While in principle this can also be achieved by skipping the non-coding exon with cryptic start codon, Hong et al excluded this possibility because in the dataset they employed, vast majority (1832/1977) of 5’ UTRs were composed of only of 1 or 2 exons. However, as we show, when one looks at all isoforms of a gene, using two current reference resources in human, vast majority (78%) of 5’ UTRs have greater than 2 exons, compelling a closer look at exon skipping as a possible mechanism underlying long first introns. Based on comprehensive analyses using multiple reference datasets, in human and in mouse, our analyses shows a strong 5’-biased exon skipping supported by a consistent bias in several genomic, contextual, epigenomic, and evolutionary features that can be functionally linked to exon skipping.

We found that the second exons often overlap with promoters or first exons of other isoforms for the same gene and this overlap is associated with a higher skipping rate of those second exons. Promoters and first exons (these are most often UTR) are known to have higher GC content than the internal exons (Zhang et al. 2004), and therefore, so do the skipped second exons. Presence of ESE motifs in an exon is associated with its inclusion (Fairbrother et al. 2002; Fairbrother et al. 2004). ESEs tend to be adenine rich (47%) and therefore a higher GC content at particular second exons results in lower density of ESE in these enhancers consistent with a higher skipping rate (Fairbrother et al. 2002; Fairbrother et al. 2004). However, in some case, higher GC-content around a splice site may promote splicing by formation of stable secondary structures (Zhang et al. 2011), and potentially counter-balance the loss of ESEs. Greater rate of skipping at second exon is consistent with an increased density of ESS, despite the fact that ESS motifs are negatively correlated with GC content, suggesting a selection to exclude these exons. Additionally, low GC-differential (relative to flanking introns) at second exon results in lower nucleosome density resulting in a greater transcription elongation rate, which is known to promote exon skipping (Amit et al. 2012).

Although, most of the investigated features mechanistically linked to exon skipping displayed 5’-bias, we did not observe any such biases for the acceptor (AS) and donor (DS) splice site scores. This is however consistent with previous evidence. Amit et al. showed that in the context of high GC differential exon, inclusion levels can lose its expected relationship with splice site score, and suggested that, higher GC differential may play a role in fine-tuning exon inclusion levels (Amit et al. 2012). Also, Shepard et al. have demonstrated that the presence of an optimal splice site does not guarantee exon inclusion (Shepard et al. 2011). In general, it is recognized that while splice sites, the polypyrimidine tract, and the branch site sequences are important, these signals alone cannot select specific splice junction, and the link between splice-site strength and splicing may not obey a simple relationship (Wang et al. 2005; Koren et al. 2007). Moreover, we note that because our exons are obtained from reference sequence data, which represent only a sample of all isoforms, rarely used exons are likely to be underrepresented in the current datasets, and it is possible that in the relatively narrow range, inclusion level is not very sensitive to splice site strength and is governed by combinatorial interactions among multiple factors. Interestingly, despite the lack of discrimination in the splice site score between the exons with different skipping rates, when we look closely at the evolutionary conservation at the most critical positions in the splice sites, namely, GT and AG, we see a lower conservation among highly skipped exons. That is, these splice sites are under a relaxed purifying constraint.

All histone marks analyzed here (except H3K36me3) have been previously linked to alternatively used exons (Table 1) (Zheng et al. 2005; Luco et al. 2010; Luco et al. 2011; Saint-Andre et al. 2011; Zhou et al. 2012; Zhou et al. 2014). For example, H3K9me3 marks at alternative exon recruit splicing factor HP1s to facilitate exon inclusion (Saint-Andre et al. 2011; Kornblihtt et al. 2013). H3K36me3 is mostly associated with constitutive exons (Kolasinska-Zwierz et al. 2009; Tilgner et al. 2009; Kornblihtt et al. 2013), however, in some instances H3K36me3 can also lead to exon skipping in collaboration with proteins MRG15 and PTB (Luco et al. 2010; Kornblihtt et al. 2013). Consistently, in our analyses second exons are enriched for such histone marks known to be associated with alternatively used exons, while H3K36me3 is enriched at downstream internal exons which tend to be constitutive.

One of our central observations is the increased rate of skipping of, predominantly UTR, exons at the 5’-end of genes. As a thought experiment, in the extreme case, certain UTR second exons may never get included in the mature transcript, thus essentially becoming cryptic or pseudo-exons (CPE). Such CPEs in some cases, we reasoned, should retain features of internal exons. This led to our expectation to observe relics of such exons in the long first introns, which we confirmed based on PIEs as a proxy for CPEs. Low GC differential at second exon is consistent with a low nucleosome density (Amit et al. 2012). However, surprisingly, PIEs have high nucleosome density despite having low GC differential. Interestingly, PIEs show high enrichment for several epigenomic marks indicative of both activating and repressive roles. Our data suggests that PIEs have regulatory roles even though they have several other features consistent with a functional exon. We speculate that over time PIEs may have acquired novel transcriptional regulatory roles, consistent with prevalent transcription factor binding at exons (Stergachis et al. 2013), as well as enrichment of regulatory epigenomic marks in long first introns (Park et al. 2014). A recent study showed that lineage-specific gained exons are alternatively spliced and preferentially localized in 5’ UTR (Merkin et al. 2015), and moreover, the novel exons exhibit higher nucleosome density, lower ESS and higher ESE (Merkin et al. 2015). Interestingly, while the putative ‘lost’ exons in our study, i.e., PIEs, also exhibit a higher nucleosome density, which we attribute to regulatory functions, PIEs show the opposite trend for ESS and ESE, consistent with their extreme skipping rate and eventual pseudogenization. We found statistical support for pseudogenization of PIEs in human (Fig. 7 and Supplementary fig. 11), based on their inclusion in cDNAs in several mammalian species. However, the fraction of PIEs detected in other species is low, which may be explained by divergent evolution of gene structure as well as sequence divergence over the long evolutionary period (Subramanian and Kumar 2003; Lorente-Galdos et al. 2013).

These findings suggest that alternative splicing toward 5’ of the gene can on one hand result in novel lineage-specific exons, while at the same time can lead to exon loss, thus impacting exon evolution in opposing manners.

Overall, our analyses offers an alternative mechanism for the underlying long first introns in higher eukaryotes, namely, that they may be a manifestation of greater skipping of 5’, mostly UTR, exons. A 5’-bias in exon skipping rate is mechanistically supported by a consistent bias in several features that are, or can reasonably be linked to exon skipping. Our estimate of exon skipping rate is based on a grouping of mRNAs by their 5’-end, which is a reasonable proxy for pre-mRNA. One can imagine that the evolutionary gain of alternative promoters in the 5’ of the gene, accompanied by a greater skipping of 5’ internal exons, may underlie the gradual increase in intron length. However, our analysis does not capture the evolutionary dynamics of the intron length increase, which would require a well-curated mRNA dataset in a large number of species. As argued, and empirically supported above, frequent skipping of exons may result in their pseudonization, that we detected as PIEs with evidence for inclusion in cDNAs in other mammals. However, interestingly, we found that PIEs are associated with weak exonic signals and evolutionary conservation, but are enriched for regulatory histone marks. While exonization of *Alu* elements in introns and new exons in 5’UTR have been observed, (Stower 2013; Zarnack et al. 2013; Merkin et al. 2015), our results suggest that in some cases, exons may be recruited to play a transcriptional regulatory role, consistent with previous reports showing enrichment of regulatory marks in long first introns (Park et al. 2014).

## Methods

### Putative pre-mRNA template preparation and filtration

The two reference exon-intron annotations for human transcripts, Ensembl (GRCh37.75) and Refseq (GRCh37/hg19) were downloaded from Ensembl and UCSC table browser respectively and removed redundancy (Karolchik et al. 2004; Cunningham et al. 2015). We only used transcripts of protein coding genes. Redundant transcripts were excluded. The rationale for our approach to construct putative pre-mRNA template is as follows. All transcripts with identical start location were considered to be results of alternative splicing of the same pre-mRNA. Therefore, the transcripts for a gene were grouped by their precise start locations and then the largest transcript-group was chosen as the representative ‘pre-mRNA’. The first exons of all transcripts in the group were collapsed to represent the first ‘meta-exon’ for the pre-mRNA. We further reasoned that any internal exon (for any mRNA for the gene) that overlapped the genomic span of the pre-mRNA, should be considered an internal exon for the pre-mRNA regardless of whether or not the exon overlaps any exon within the pre-mRNA group. Thus to define the internal exons of the meta-transcript, we collapsed all internal exons for all the transcripts of the gene into internal meta-exons. The exon collapsing was done based on genomic location overlap (Fig. 1). Only the meta-transcripts having greater than four meta-exons were retained. This procedure yielded 7,205 putative pre-mRNA templates. From these templates, we only used internal exons having length (50bp-300bp) and flanking long introns (> 300bp) for further investigation. Also, the 74 putative pre-mRNA templates that had redundant meta-exons and pure introns relative to another pre-mRNA template were excluded from further analysis.

### Estimating exon skipping rates

For the reference dataset above, we obtained fraction of transcripts (within a group) where a particular base of a meta-exon was included in the transcripts. The average base-level inclusion was used as the exon-level inclusion level. For exon skipping rate for the HEXEvent (Busch and Hertel 2013), we downloaded exons inclusion levels, and transformed the inclusion level into a skipping rate (1 minus inclusion rate). MISO database (Katz et al. 2010) provides the coordinates of skipped exons. These annotations were mapped to our putative pre-mRNA templates. For each ordinal position (say the second exon), we estimated the fraction of all second exons in the genome that were skipped according to MISO, and compared this quantity for different ordinal positions.

### Obtaining, calculating and mapping the features

The sequences of acceptor and donor splice site are obtained from only internal exons and the scores of those were calculated by MaxEntScan (Yeo and Burge 2004). The scores of multiple alternative splice sites in each exon were averaged. The proportions of ESE and ESS in exons are computed by sliding window using the hexamers reported in RESCUE-ESE (Fairbrother et al. 2004) and FAS-ESS (Wang et al. 2004). The phastCons scores for 100 species and 46 species subsets were downloaded from the UCSC and then mapped into putative pre-mRNA templates (Karolchik et al. 2004; Siepel et al. 2005). Evolutionary conservation of each exons is calculated as mean of phastCons score on their exons. All signals (density graph) as a epigenomic feature including nucleosome density, H3K36me3, H3K4me1, H3K27ac, H3K9me3 and H3K27me3 measured in GM12878 and K562 were downloaded from the “Histone Modifications by ChIP-seq from ENCODE/Broad Institute” download page of the UCSC (Kent et al. 2002; Bernstein et al. 2005; Bernstein et al. 2006; Mikkelsen et al. 2007; Guttman et al. 2010; Ernst et al. 2011; Valouev et al. 2011). All signals were mapped into all internal exons of putative pre-mRNA templates and flanking 200bp intron region. When summarizing an epigenomic feature along the exon body, to accommodate the variation in exon lengths, each exon was segmented into 50 equal-sized bins and signal obtained for each bin.

### Putative Internal Exons (PIE)

From the putative pre-mRNA templates, we obtained the pure introns, which do not have any internal exons of protein coding mRNA. We used GENSCAN (Burge and Karlin 1997) to predict PIEs in pure introns and excluded short predicted PIEs (< 50bp). From the predicted PIEs, we excluded the PIEs overlapping with other overlapping genes (e.g., the second exon of MYZAP gene (ENSG00000263155) is in third pure intron of GCOM1 (ENSG00000137878)), the first and last exons of other mRNAs which are not included in generating the putative pre-mRNA, the pseudogenes, the noncoding RNAs and the PIEs harboring repeat element predicted by RepeatMasker (Smit et al. 2013-2015).

### BLAST search of PIEs against 37 mammals

We obtained cDNA sequences (GRCh37.p13) of 37 mammals from the Ensemble (ensembl.org). The species-specific repeats and low-complexity sequences on the cDNA sequences were masked by RepeatMasker (Smit et al. 2013-2015) and Dustmasker of NCBI-BLAST-2.2.31+ (Altschul et al. 1990; Camacho et al. 2009). We used the BLASTn program to search for PIEs with megablast algorithm (word size = 11) against the 37 mammals masked cDNA databases.

## Acknowledgments

This work was funded by R01 GM100335 to S.H. Authors would like to thank Dr. Steve Mount for helpful discussions, and Shrutii Sarda and Dr. Nishanth Nair for comments on the draft.

## Author contributions

SGP conceived the initial idea and developed the analyses with help from S.H. SGP performed all analyses. SH and SGP interpreted the results and wrote the manuscript.

## Disclosure declaration

None

